# Whole mitochondrial genome sequencing- a novel approach for studying Phylogenomics for *Anas platyrynchos*

**DOI:** 10.1101/2022.07.11.499548

**Authors:** Aruna Pal, Argha Chakraborty, Manti Debnath

## Abstract

The evolution of duck and its distribution across the globe has been an interesting aspect. Domestication of duck and its evolution was also important to analyze if domesticated duck has migrated to other region of the globe or separate domestication process has occurred in particular region of the world. Mitochondrial genes as D-loop or cytochrome B had been used for studying evolution or molecular phylogeny. In current study, we aim to study phylogenomics for *Anas platyrynchos* across the globe. Whole mitochondrial genome sequencing for Bengal duck (indigenous duck from West Bengal, India) was performed on illumina platform and gene sequences were analyzed for all 37 mitochondrial genes. Variabilities were observed at nucleotide sequence, but absent in amino acid sequence and corresponding 3D structure of polypeptides coded by mitochondrial genes. We observed *Anas platyrynchos* from Eurasia (europe and Asia) were clustered together indicating common origin. Extensive variabilities were observed among the chinese duck population. The duck population from US was outgrouped. These ducks might have migrated from China and later bred with wild ducks to evolve as new breed or strain in US.

## Introduction

Domestic duck (*Anas platyrynchos*) is mostly distributed world wide and has the potentiality to evolve as leader in poultry industry. In the current time, ducks have acquired almost second position as poultry. Traditionally ducks are reared in waterbodies, occurs in shallow freshwater beels, marshes, reservoirs, lakes, rarely rivers and ponds., but due to gradual advancement of management practices, they can be reared in intensive management practices. Ducks are reared as layer and broiler depending upon the breed and the choice of the farmers. Ducks were observed to have the unique properties of better resistance among the avian diseases compared to chicken, which has gained its popularity^**1,2,3**^. We have explored the genomics for duck in terms of innate immune response genes^**4-10**^. We have explored duck immune response genes against duck plague infection^**11-12**^.

It has become exceedingly important to study the biodiversity as well as evolutionary significance of duck. Reports indicate domesticated ducks evolve from the wild relatives^**13**^. Ducks initially originate from Africa (Nile Valley), South Eastern Asia (Indus Valley to South Eastern China), Mexico and Cuba, later ‘introduced to South Africa, Mauritius, Australia and New Zealand, among others. Although their distribution is spread over the entire world, they are dominated in northern hemisphere^**14**^.

Certain studies have been undertaken to understand the molecular evolution and studies with mitochondrial gene was found to be promising. Most of the studies have been observed for D loop, cytochrome B and ND2 gene among the mitochondrial genes^**15-20**^. Phylogenetic analysis for Bengal duck (indigenous duck for West Bengal, India) have been studied in our lab among the duck population reared in different agroclimatic zone of West Bengal based on biomorphometric characteristics^**22, 1**^. Reports are available for phylogeny and evolutionary ecology of Modern sea ducks^**23**^.

We have sequenced cytochrome B, cytochrome C and D loop (control region) among the mitochondrial genes for these Bengal ducks from five different agroclimatic conditions of West Bengal, duck population were found to be absent from hilly region (Project report, SERB).

## Materials and methods

### Whole mitochondrial genome sequencing

Whole mitochondrial genome analysis (WMGA) was conducted for Bengal duck from West Bengal of India (Genebank accession number MN011574). The mitochondrial map was generated. The steps for next generation sequencing studies (Whole mitochondrial genome sequencing) involves isolation of mtDNA, qualitative and quantitative analysis of g-DNA: PCR amplified with COX-2(mt specific), GAPDH and Beta actin primers to validate. The next step involves preparation of library: The paired-end sequencing library (NEBNext Ultra DNA Library Preparation Kit), quantity and quality check (QC) of library on Bioanalyzer: Bioanalyzer 2100 (Agilent Technologies) using High Sensitivity (HS) DNA chip. The final step is cluster Generation and Sequencing. The adapters are designed so as to allow selective cleavage.

Whole mitochondrial genome sequences for Anas platyrynchos were retrieved from other parts of the world. However the geographical distribution for the published complete mitochondrial sequences were not uniform. Most of the sequences were available from China, Asia, however single sequence (MN720361) from Norway, Europe, and another sequence from USA (NC_009684), North America

### Construction of Phylogenetic tree

Phylogenetic tree was constructed based on neighbour joining method with Jukes-Cantor substitution model of MAAFT software^**24**^ with all the available sequences from gene bank. Till date complete mitochondrial sequences are available for duck (Anas platyrynchos) from China, India, Norway and US. Extensive variability was observed among the duck population of China.

We generated phylogenetic tree among the duck population of these countries. The sequence genetically closest to the consensus sequence was taken for consideration in case of Chinese duck.

### Comparison of mitochondrial gene sequences for Indian duck (Bengal duck) with ducks from other parts of the world

Nucleotide sequences such otained was subjected to multiple sequence alignment MAFFTsequences. Alignment report was visualized in MSA viewer. SNPs were identified through Lasergene DNASTAR software^**25**^.

### Comparative analysis for amino acid sequences for the coding genes for Indian duck with respect to others

Derieved amino acid sequences for all the 13 coding genes for mitochondrial genes were aligned for the detection of amino acid sequence variabilities.

### Three-dimensional structure prediction and Model quality assessment

Accordingly in the next step, 3D structural analysis for the polypeptide (13 in numbers) were predicted. The templates which possessed the highest sequence id entity with our target template were identified by using PSI-BLAST (http://blast.ncbi.nlm.nih.gov/Blast). The homology modeling was used to build a 3D structure based on homologous template structures using PHYRE2 server^**26**^. The 3D structures were visualized by PyMOL (http://www.pymol.org/) which is an open-source molecular visualization tool. Subsequently, the mutant model was generated using PyMoL tool. The Swiss PDB Viewer was employed for controlling energy minimization. The structural evaluation along with a stereochemical quality assessment of predicted model was carried out by using the SAVES (Structural Analysis and Verification Server), which is an integrated server (http://nihserver.mbi.ucla.edu/SAVES/). The ProSA (Protein Structure Analysis) webserver (https://prosa.services.came.sbg.ac.at/prosa) was used for refinement and validation of protein structure^**27**^. The ProSA was used for checking model structural quality with potential errors and the program shows a plot of its residue energies and Z-scores which determine the overall quality of the model. The solvent accessibility surface area of the genes was generated by using NetSurfP server (http://www.cbs.dtu.dk/services/NetSurfP/)^**28**^. It calculates relative surface accessibility, Z-fit score, the probability for Alpha-Helix, probability for beta-strand and coil score, etc. TM align software was used for the alignment of 3 D structure of IR protein for different species and RMSD estimation to assess the structural differentiation^**29**^. The I-mutant analysis was conducted for mutations detected to assess the thermodynamic stability. Provean analysis was conducted to assess the deleterious nature of the mutant amino acid. PDB structure for 3D structural prediction of gene for duck was carried out through PHYRE software^**26**^. Protein-protein interaction have been studied through String analysis^**30**^, along with the details of the bioinformatics software^**31**^.

## Result

### Characterization of whole mitochondrial genome for Bengal duck from India

WMGA was conducted for Bengal duck from West Bengal of India (MN011574). The mitochondrial map generated is being represented in Fig 1. Thirteen polypeptide coding genes were identified with their resective protein ids being <u>QVX28256.1</u> (ND1), <u>QVX28257.1</u>(ND2), <u>QVX28258.1</u>(COX1), <u>QVX28259.1</u>(COX2), <u>QVX28260.1</u> (ATP8), <u>QVX28261.1</u> (ATP6), <u>QVX28262.1</u> (COX3), <u>QVX28263.1</u> (ND3), <u>QVX28264.1</u> (ND4L), <u>QVX28265.1</u> (ND4), <u>QVX28266.1</u> (ND5), <u>QVX28267.1</u> (CYTB,), <u>QVX28268.1</u> (ND6).

**Fig 1:**
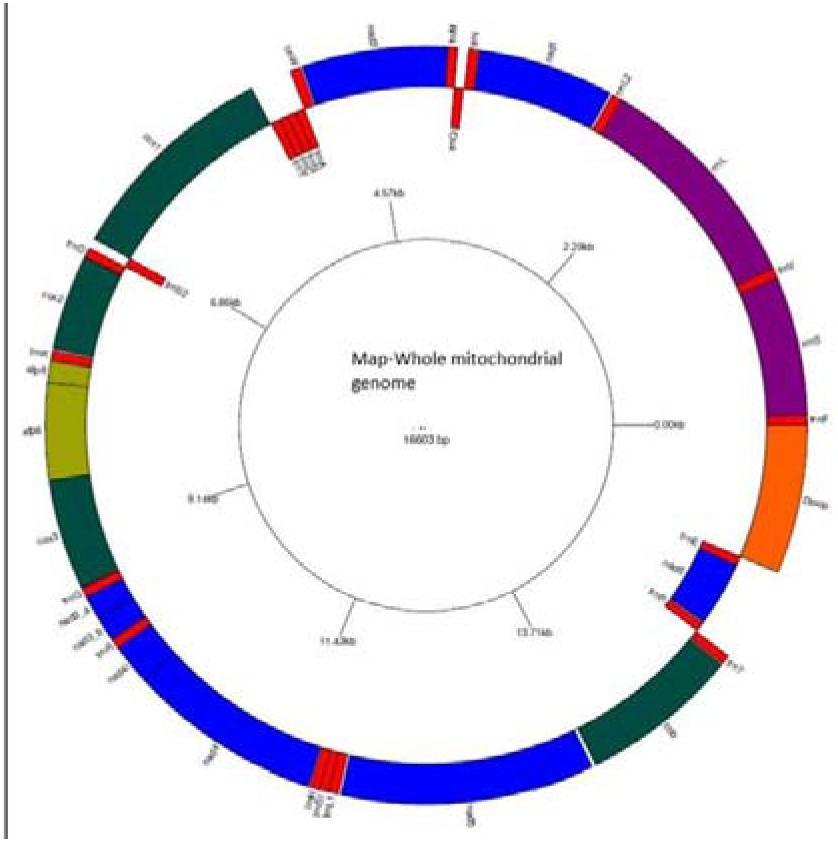
Whole mitochondrial map for *Anas platyrynchos* from West Bengal, India.

Whole mitochondrial genome sequences for ducks were retrieved from other parts of the world. However the geographical distribution for the published complete mitochondrial sequences are not uniform. Most of the sequences were available from China, Asia, however a single sequence (MN720361) from Norway, Europe, USA & China (NC_009684),North America (same sequence submitted from China also).

### Construction of Phylogenetic tree

Phylogenetic tree was constructed based on neighbour joining method with Jukes-Cantor substitution model of MAAFT software with all the available sequences from gene bank as represented in Fig 2. Till date complete mitochondrial sequences are available for duck (Anas platyrynchos) from China, India, Norway and US. Extensive variability was observed among the duck population of China.

**Fig 2:**
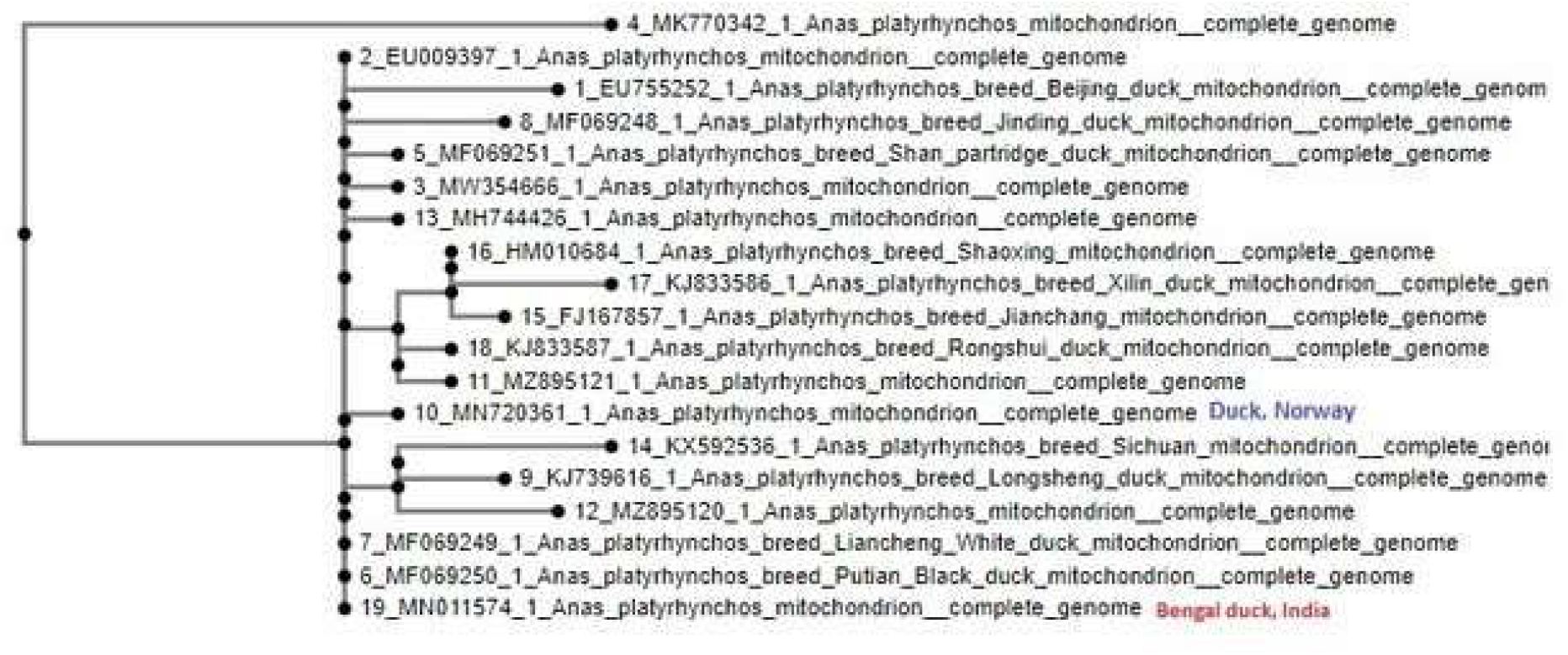
Molecular phylogeny for *Anas platyrynchos* across the globe with whole mitochondrial genome sequencing.

We generated phylogenetic tree among the duck population of these countries (Fig 3). The sequence genetically closest to the consensus sequence was taken for consideration in case of Chinese duck.

**Fig 3:**
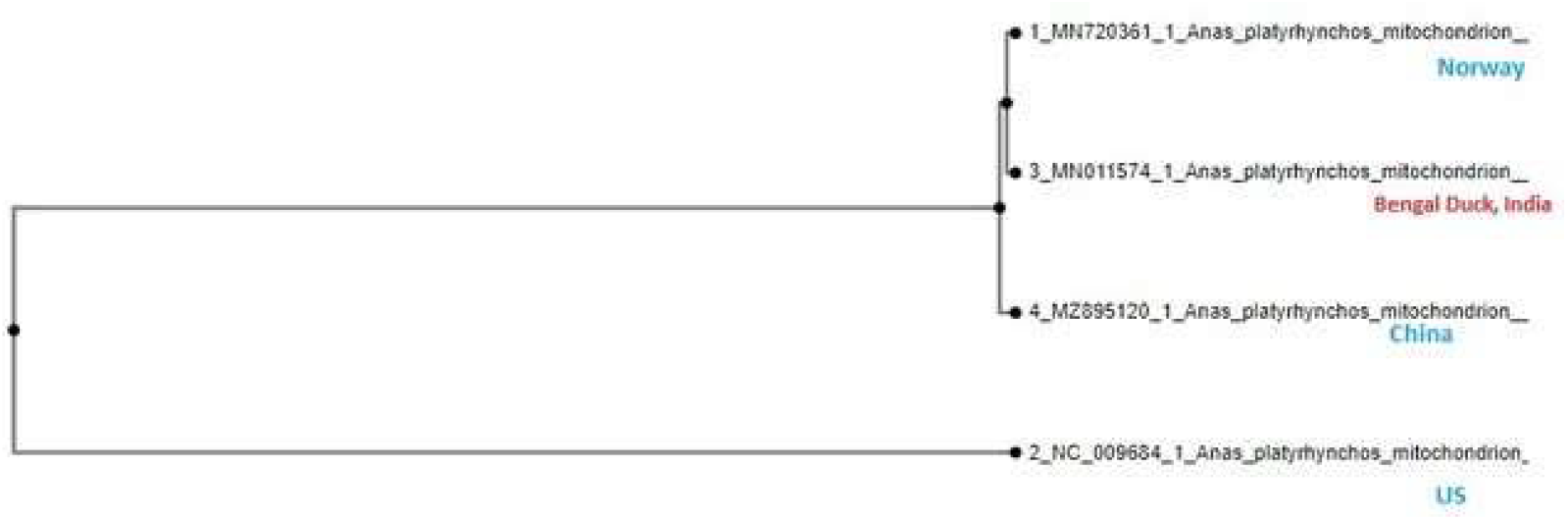
Phylogenomics through countries with whole mitochondrial genome sequencing

### Comparison of mitochondrial gene sequences for Indian duck (Bengal duck) with ducks from other parts of the world

The complete mitochondrial genome sequence for Indian duck was compared with that of mitochondrial sequences of duck from China, Norway and US (Fig 4). Certain SNPs were identified among the sequences. But the most interesting fact to note is that the mutations were non-synonymous without amino acid differences detected. We studied amino acid sequence for ND5 across different countries and amino acid identity was detected (Fig 5), irrespective of nucleotide sequence variabilities.

**Fig 4:**
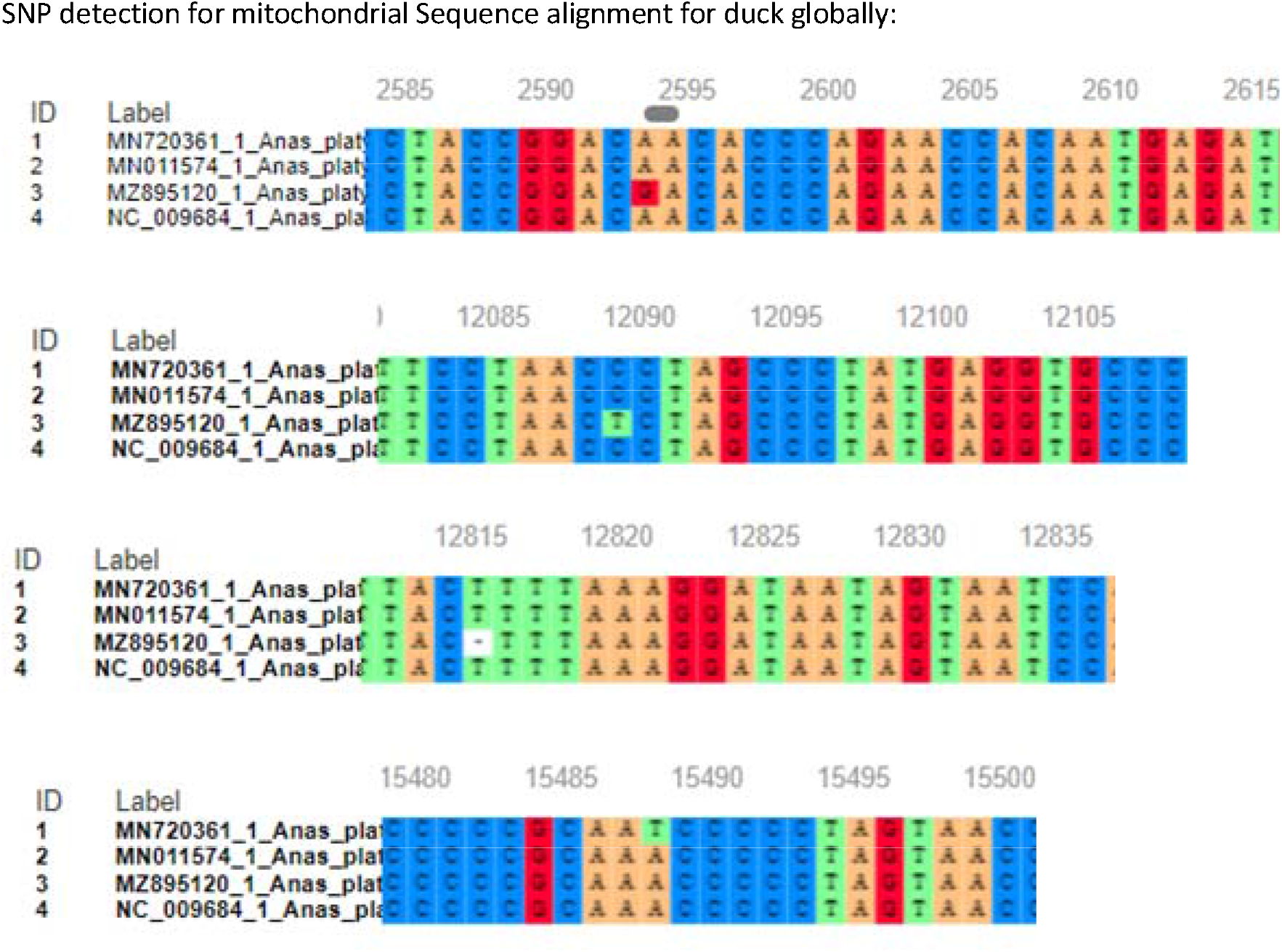
SNP detection across the mitochondrial genes for Anas platyrynchos from different countries

**Fig 5:**
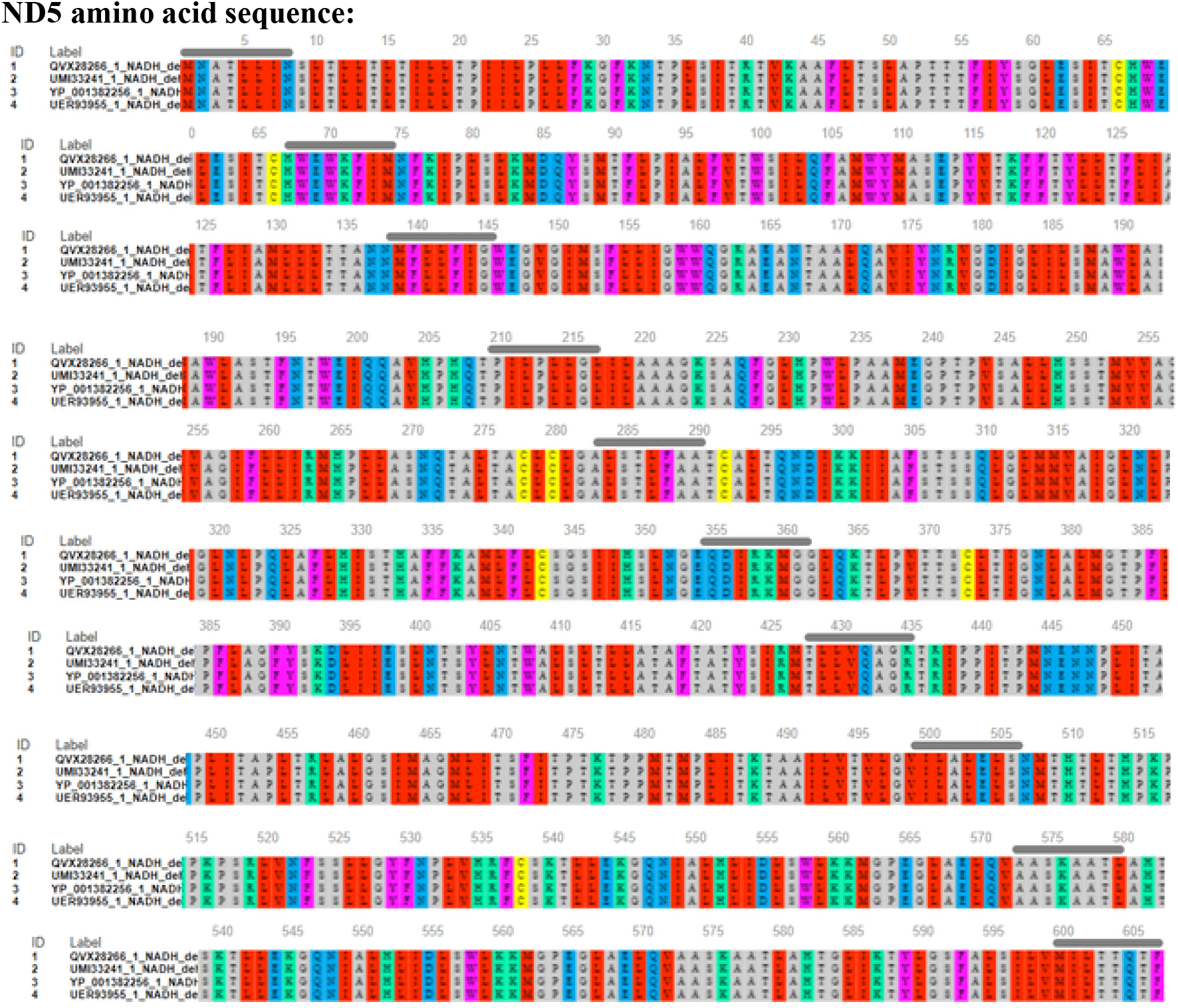
Amino acid sequence identity detected among the ND5 sequence of Indian duck (Bengal duck) with the duck from other parts of the World.

### Comparative analysis for amino acid sequences for the coding genes for Indian duck with respect to others

Out of total 37 genes, 13 genes codes for polypeptide. Derieved amino acid sequences for these genes were compared for ND5 to reveal indentity (Fig 5). The molecular interaction for mitochondrial gene has been depicted in figure 6

**Fig 6:**
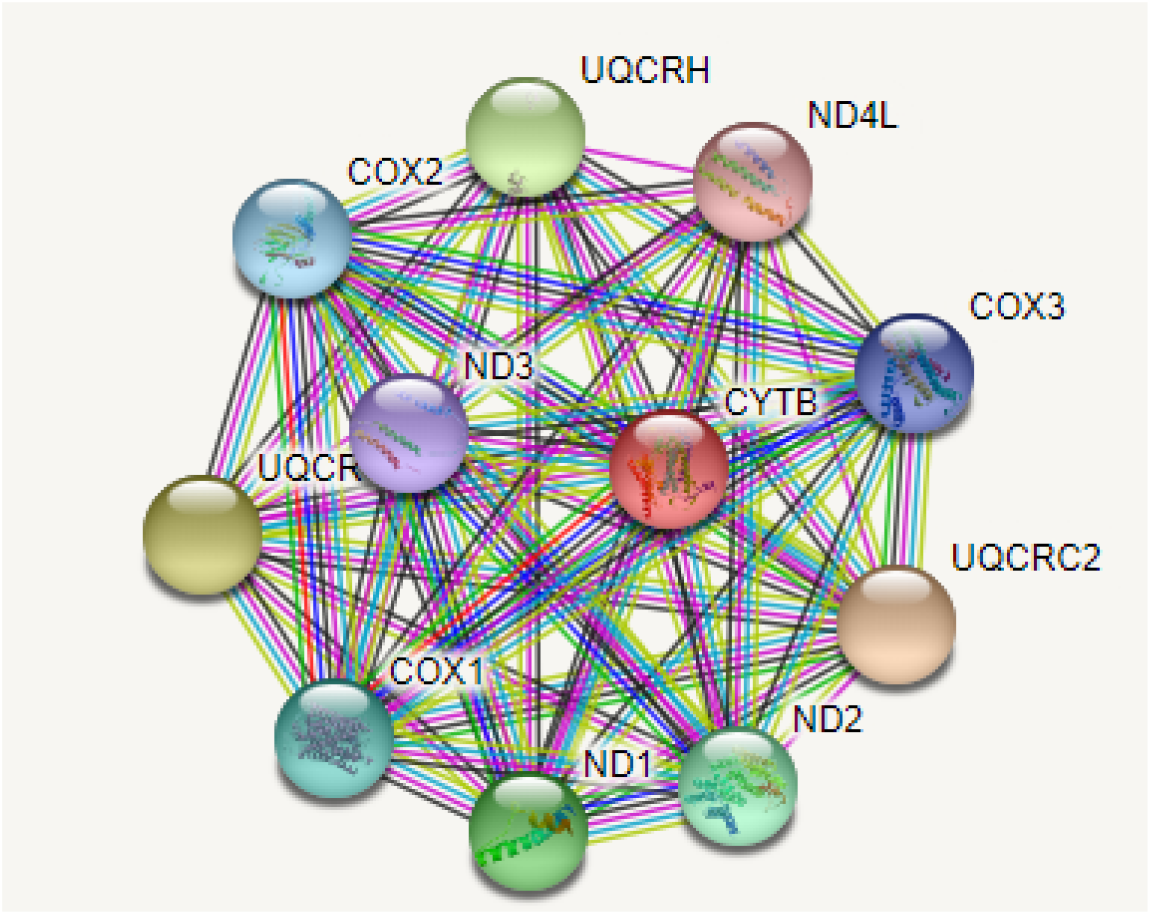
Molecular interaction studies among the Anas platyrynchos genes through String analysis

### 3D structural analysis for derived peptides for the coding genes of mitochondria

The whole mitochondrial genome sequencing `MN011574)` reveals 37 genes with 13 coding genes, non coding genes and control D-loop region. 3D structural analysis of derieved amino acids with respect to 13 polypeptide genes were conducted. Although sequence variabilities were observed at nucleotide level, aminoacid sequenceforthese 13 coding genes for mitochondria were observed to be conserved across the globe. Alignment for the 3D structure for these coding genes were found to have RMSD as zero, implying no structural variation (Fig7).

**Fig 7:**
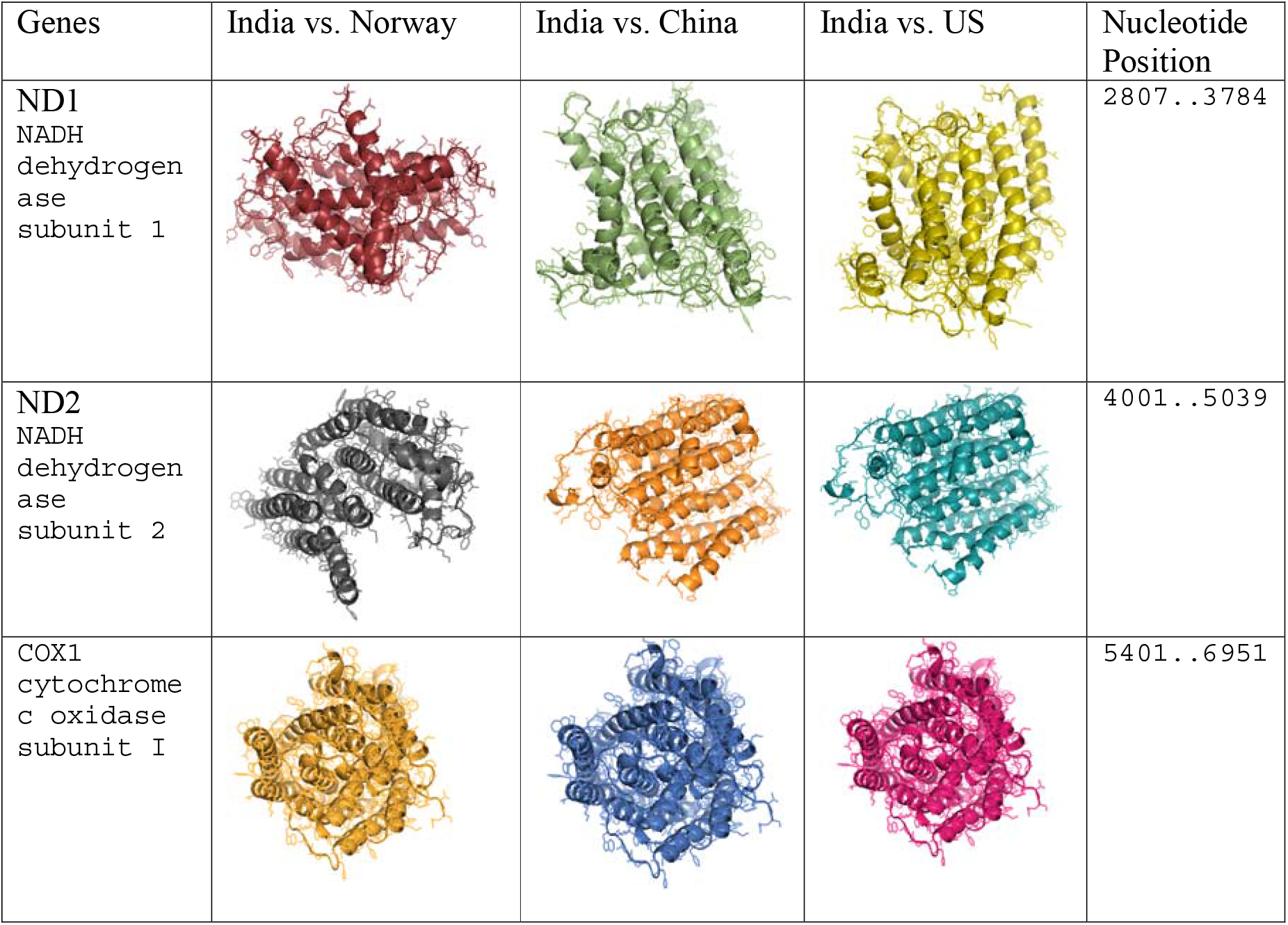
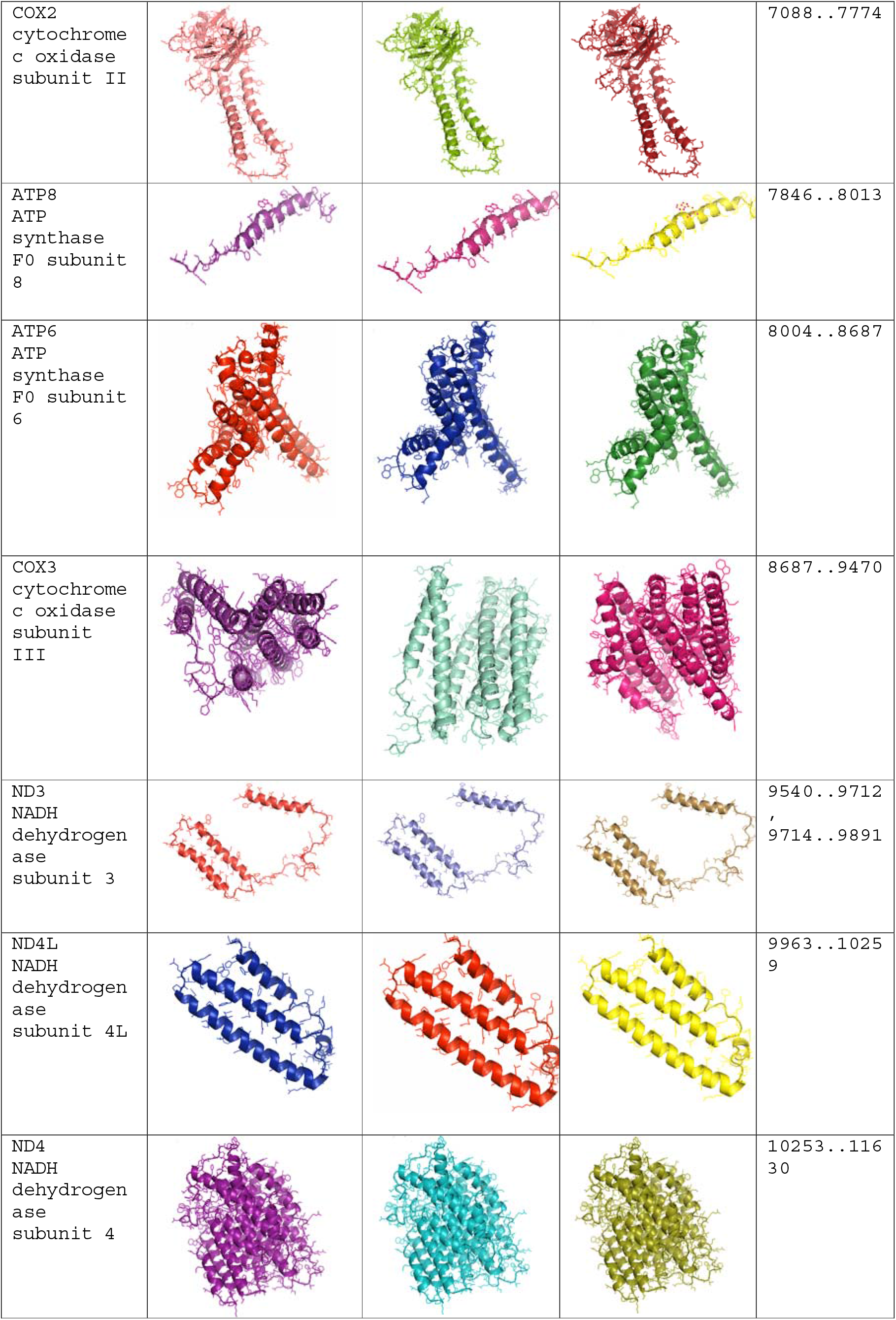
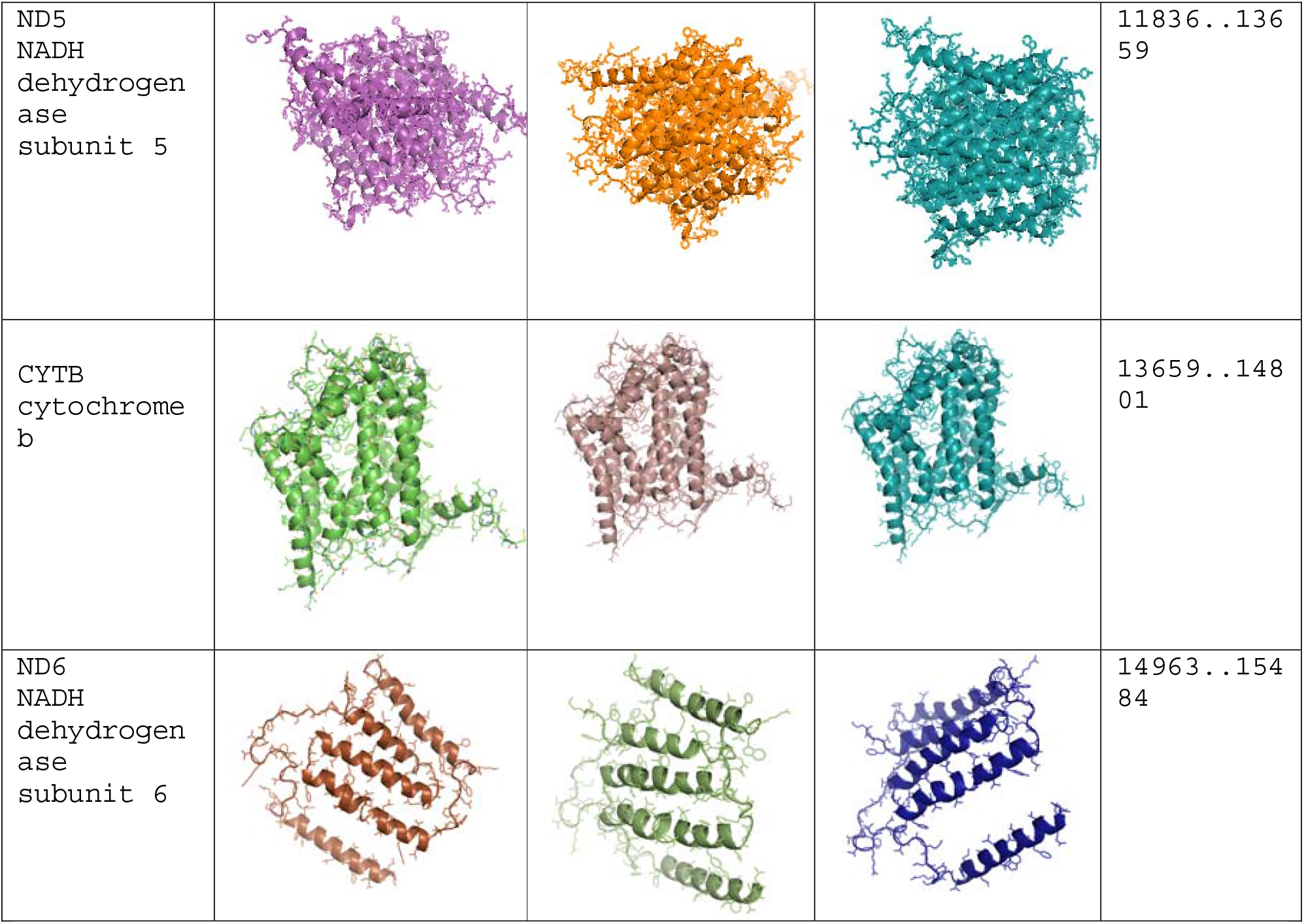
Alignment of the derived peptide from coding sequence of mitochondrial gene (3D structure) in Indian duck compared to other ducks distributed globally.

## Discussion

Ducks (Anas platyrynchos) have evolved time immorial across the globe. Reports are available that domesticated ducks that we rear today have originally evolved from wild ducks. A class of migratory ducks are available, which may be a source of global distribution. The natural habitat for migratory ducks are shallow or marshy water bodies. The domesticated ducks are mostly reared for its nutritious eggs providing nutritional security to the rural woman and children. It also acts as economic insurance to the rural community or farmers. Duck meat is also a delicacy, specially in winter. It is rich in certain class of nutrients and comparable to mutton or chevon. The major important characteristics of indigenous duck is that they are mostly resistant to common avian diseases.

It has been a curiosity to study the evolution or domestication of ducks. Another major concern for duck is that they were observed to be asymptomatic to avian influenza^**2**^. It has equally been claimed that wild migratory birds act as major source for spread of avian influenza virus. Domesticated ducks that swim in waterbodies nearby the village act as a carrier of the virus and spread them within the chicken population in the village. Evolution of ducks has been studied by a number of scientists across the globe based on D-loop, ND or Cytochrome B gene as mitochondrial gene^**15-20**^. The current study revealed that mitochondrial genes with all 37 genes including 13 coding genes and rest non coding and a control region (D loop), provide a better in site for evolutionary studies. The current study is unique in the sense that it employs all 37 mitochondrial genes for studying evolutionary studies for Anas platyrynchos across the globe, instead of few genes for the first time.

In the current study, we had analyzed whole mitochondrial genome sequencing for the indigenous ducks (Bengal duck). The mitochondrial genome consists of 13 protein-coding genes, 2 rRNA genes (12S rRNA and 16S rRNA), 22 tRNA genes, and 1 control region (CR). The total length of the genome is 16,623⍰bp, with a base composition of 29.66% A, 22.28% T, 15.35% G, and 32.71% C. The total length of 13 protein-coding genes is 10,997⍰bp long, all of which are encoded on the same strand except for ND6 in the light strand. Except for ND2, ND5 and Cytb (ATC start codon), ND6 (TAA start codon), and COX1,COX2 (GTG start codon), the remaining 7 protein-coding genes initiate with ATG (COX3, ND1, ND3, ND4, ND4L, ATP6, ATP8). and retrieved the gene bank sequences for complete mitochondrial genome from *Anas platyrynchos* across the globe. Mitochondrial genome map comprises of 37 genes with 13 polypeptide coding genes. We had also observed from string analysis that there is interaction among these mitochondrial genes. As a result the comprehensive study involving all mitochondrial genes provides us with better research output. We could retrieve such complete mitochondrial sequences from four countries only, namely India, China, Norway and US. Wide variabilities have been observed among the chinese duck population. A complete outgroup was observed from China as accession no. MK770342.

In order to have a better understanding, we use the chinese duck sequence which is closest to the consencus sequence among the duck population from China. The lead to a better visualization of the phylogenetic tree. Indian duck population was observed to genetically closest to that of duck from Norway. Ducks from Eurasia were grouped together.

Earlier observations have also indicated ducks initially originated from Asian and Europen countries. Later on some ducks like Pekin were transported to US, where they bred to local duck and current day US duck population evolve.

A group of scientists from have studied phylogenetic relationships among the mitochondrial gene as Cytochrome B among east asian ducks. They observed Japanese and Chinese ducks were clustered to the same clade distinct from the clade containing European and Southeast Asian ducks. domesticated ducks in China and Southeast Asian countries have been distributed to Japan and European countries, respectively. In addition, this phylogenetic tree revealed that the genetic constitution of Southeast Asian ducks is distinctly different from that of Northeast Asian ducks^**15**^.

Similarly the variability in mitochondrial cytochrome C oxidase I was also employed for studying phylogenetic analysis for ducks in Indonesia^**34**^. Similarly, D loop sequences or control region is used for studyion phylogenetic relationships among the duck population of Nigeria^**16**^. Certain reports are available for studying evolution through mitochondrial genes^**32-34**^.

Comparison of the nucleotide sequences have revealed certain SNPs among the mitochondrial genes. It has been interesting to note that the SNPs don’t lead to any variation in the amino acid sequence, while alignment for derived amino acids from the thirteen numbers of mitochondrial polypeptides were studied. 3D structural prediction for the mitochondrial polypeptides were studied for ducks from India, Norway, China and US. Structural alignment for the 3D structures for these mitochondrial proteins reveal RMSD value as zero, indicating structural similarities. This indicates the structural uniqueness of the mitochondrial proteins, which play a major role in oxidative phosphorylation and other essential vital role of the body. Thus whole mitochondrial genome sequence may evolve as a promising tool for studying duck evolution, which had added importance for zoonotic importance.

## Acknowledgement

The authors are thankful to Department of Biotechnology, Ministry of Science and Technology, Govt. of India (Grant number BT/PR24310/NER/95/649/2017) and Science and engineering research Board, Department of Science and Technology, Govt. of India (Grant no. EMR/2016/003554) for providing the financial support. The technical and financial support by Vice-Chancellor, West Bengal University of Animal and Fishery Sciences is duly acknowledged. Thanks to Director, AH & VS, Animal Resource Development Department, Govt. of West Bengal.

